# Circularly permuted LOV domain as an engineering module for optogenetic tools

**DOI:** 10.1101/2021.05.19.444876

**Authors:** Lequn Geng, Jiaqi Shen, Wenjing Wang

**Affiliations:** Life Sciences Institute, University of Michigan, Ann Arbor, Michigan; Department of Chemistry, University of Michigan, Ann Arbor, Michigan

## Abstract

We re-engineered a commonly-used light-sensing protein, LOV domain, using a circular permutation strategy to allow photoswitchable control of the C-terminus of a peptide. We demonstrate that the use of circularly permuted LOV domain on its own or together with the original LOV could expand the engineering capabilities of optogenetic tools.

## Main Text

Optogenetic tools have been transformative by enabling manipulation of specific cellular processes using light^1–3^. Their genetic encodability and light-dependence allow fast and reversible control of cellular events in specific cell types. Light-sensing proteins^4^ are crucial building blocks for engineering optogenetic tools. Among them, the second light, oxygen, voltage sensing domain from *Avena sativa* phototropin 1 (AsLOV2) has been most well-studied^5–8^ and most widely applied to modulate the activity of various proteins and peptides^9–23^.

The broad applicability of AsLOV2 is due to its unique mechanism for light-dependence. In the dark state of AsLOV2, its C-terminal helix, termed Jα-helix, is packed against the Per-Arnt-Sim domain (PAS domain, hereafter referred to as the “protein core”) through hydrophobic packing and hydrogen bonding interactions^5^ (Fig. 1A). With light irradiation, a cysteine residue in the protein core forms a covalent bond with the cofactor flavin mononucleotide, and the structural change propagates to the Jα-helix, causing it to unwind from the rest of the protein^5^. The main approach to render a peptide or protein photoswitch is therefore to directly fuse it to the C-terminus of the Jα-helix. This introduces steric hindrance (“blocking”) in the dark state which is “unblocked” in the light state.

**Figure 1:**
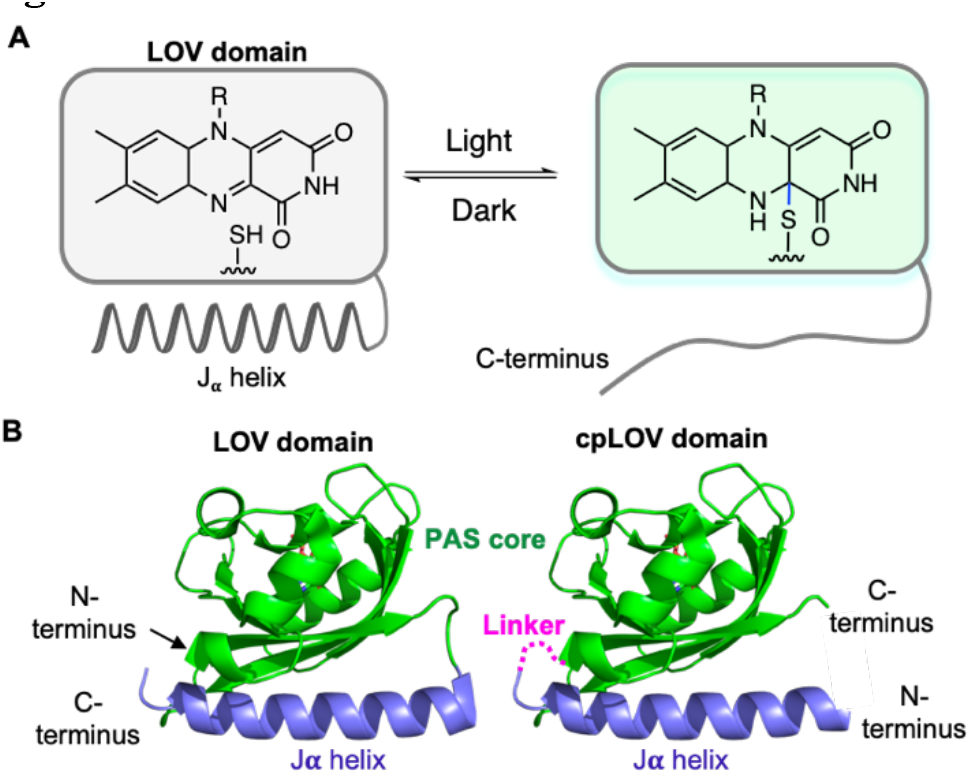
(A) Light-dependent reversible conversion of the LOV domain. (B) Design of the circular permuted LOV (cpLOV) based on the LOV domain (PDB: 2V1A).

While this light-dependent conformational change of AsLOV2 has been useful for protein engineering, it limits the use of AsLOV2 to only cage the N-terminus of a peptide and precludes the possibilities to cage peptides where their function requires a free N-terminus. In addition, for some peptides, it will be more effective to cage the C-terminus that contains critical residues. Therefore, a similar light-sensing domain that can cage the C-terminus of a peptide will complement AsLOV2 and expand the designs of optogenetic tools.

Herein, we report a re-engineered AsLOV2 domain using a circular permutation strategy (cpLOV). We demonstrate that cpLOV retains light-sensing capability while allowing modulation of peptides’ activities by controlling their C-terminus. In contrast to a conceptually similar design^24^ published recently, our cpLOV allows peptides to have a free N-terminus. In addition, we show that simultaneous caging of both the N- and C-termini of a peptide using cpLOV and AsLOV2 together (hereafter referred to as “dual caging”) provides enhanced caging of the peptide target and alters the dynamic range of an existing optogenetic tool. Therefore, cpLOV represents a new light-sensing module useful for engineering optogenetic tools.

To engineer cpLOV, we started from an AsLOV2 variant, hLOV1^25^, which contains 15 mutations from the wild-type AsLOV2 and has been shown to have superior caging in the dark state. We first connected the hLOV1 termini with a flexible linker. Based on the crystal structure^5^ (Fig. 1B), we reasoned that a four-amino-acid (GSGS) linker is sufficient to connect the original N- and C-termini of the LOV domain. We then introduced a new opening at the original “hinge region” connecting Jα-helix to the protein core. We split between amino acids L520 and H521, as H521 is the first helical residue on the Jα-helix.

To check whether the light-induced conformational change of Jα-helix could still take place in cpLOV, we used cpLOV to cage a heptapeptide, SsrA, and used a yeast surface-based binding assay with SspB protein (Fig. 2A) to evaluate its accessibility in the dark and light states. Yeast surface display was used because this allows future directed evolution^26,27^ to improve cpLOV. We screened 10 fusion sites along the Jα-helix (Fig. 2B & C), from V529 to Q538, as it has been shown that the fusion sites on the Jα-helix affect the light-dependence of each peptide^13,19,23^. Four out of the ten constructs tested showed light-induced signal increase (Fig. 2D & 2E). The best-performing construct, cpLOV(b), showed higher background in the dark but also higher signal in the light compared to the hLOV1-SsrA fusion protein. This shows that cpLOV has light-sensing capability. Interestingly, the four constructs with light dependence were fused to the two ends of the Jα-helix, away from the middle section of the Jα-helix. This extensive study of the cpLOV fusion constructs suggested the importance of testing various fusion site on the Jα-helix to achieve an optimal light-dependence.

**Figure 2:**
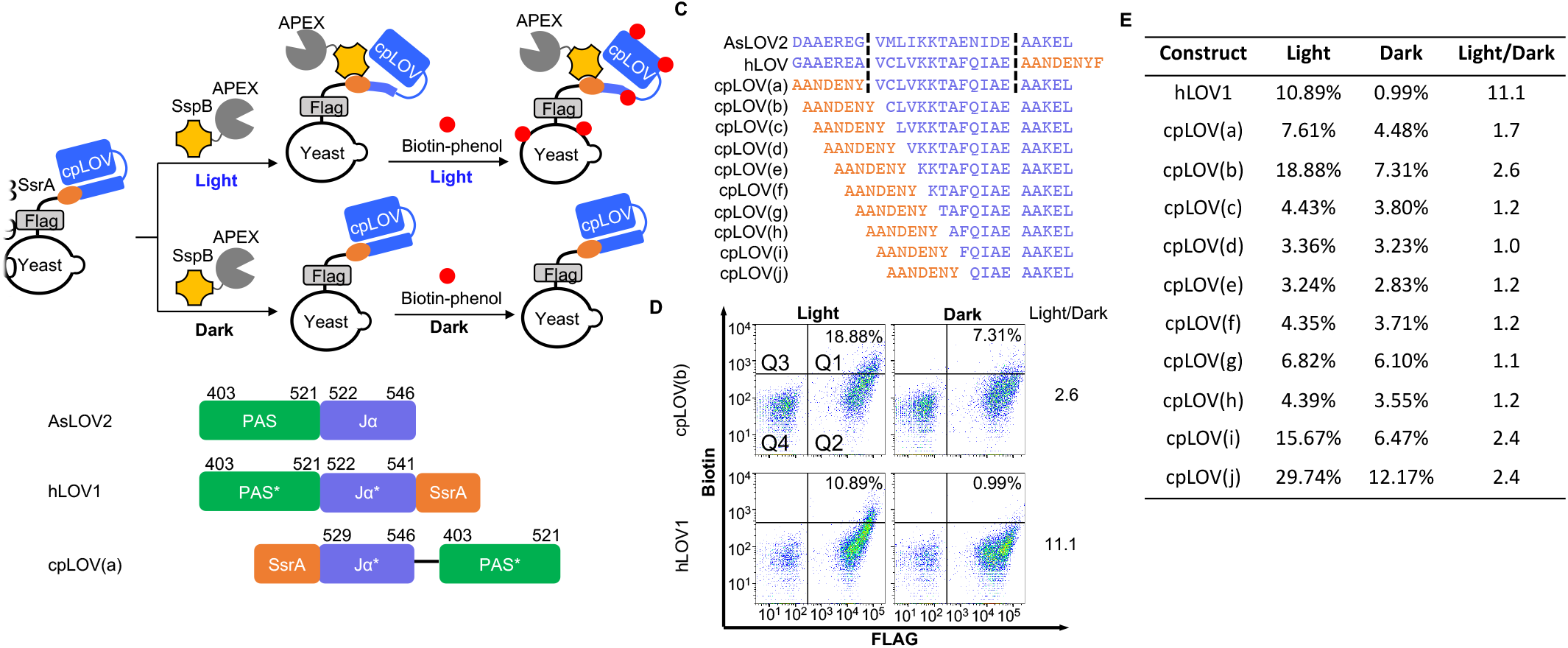
(A) Yeast surface display assay for testing cpLOV’s caging of SsrA peptide. cpLOV is expressed on the yeast surface. Under light irradiation, SsrA is uncaged and recruits SspB-APEX fusion protein. The APEX can covalently label proteins in proximity with biotin-phenol molecule. (B) The architecture of AsLOV2, hLOV, and cpLOV(a) domain. In cpLOV(a), the original N- and C-termini are linked through a four amino acid GSGS linker. (C) The Jα helix sequences and truncation sites for different cpLOVs. SsrA peptide sequence, AANDENY. The SsrA sequence for hLOV is adapted from the previously reported iLID23. (D) Flow cytometry analysis of the best SsrA-cpLOV fusion construct. The quadrant was drawn based on the negative population in the bottom left corner in Q4. The percentage in Q1 indicates the ratio of the cell count in Q1 to that in (Q1 + Q2). The Light/Dark value is the ratio of the percentage of the light to that of the dark state. (E) Light response of cpLOV constructs.

We then asked whether cpLOV can be generally applied to cage other short peptides of chemical and biological interests. We chose a heptapeptide, tobacco etch virus protease cleavage site (TEVcs), as TEVcs has been widely used in synthetic biology^25,28–30^ due to its specificity and orthogonality in eukaryotic cells. Using the yeast surface display platform (Fig. 3A), we tested the light-dependent protease cleavage of a TEVcs-cpLOV fusion construct (Fig. 3B). As shown in Fig. 3C, the best fusion construct showed a light dependent protease cleavage in between the AsLOV2 and hLOV1 domains. Even when used in a highly-sensitive transcriptional assay in HEK293T cell culture (Fig. 4A), cpLOV also provided sufficient caging in the dark and resulted in 5.6-fold light-dependent transcriptional activation (Fig. 4B).

**Figure 3:**
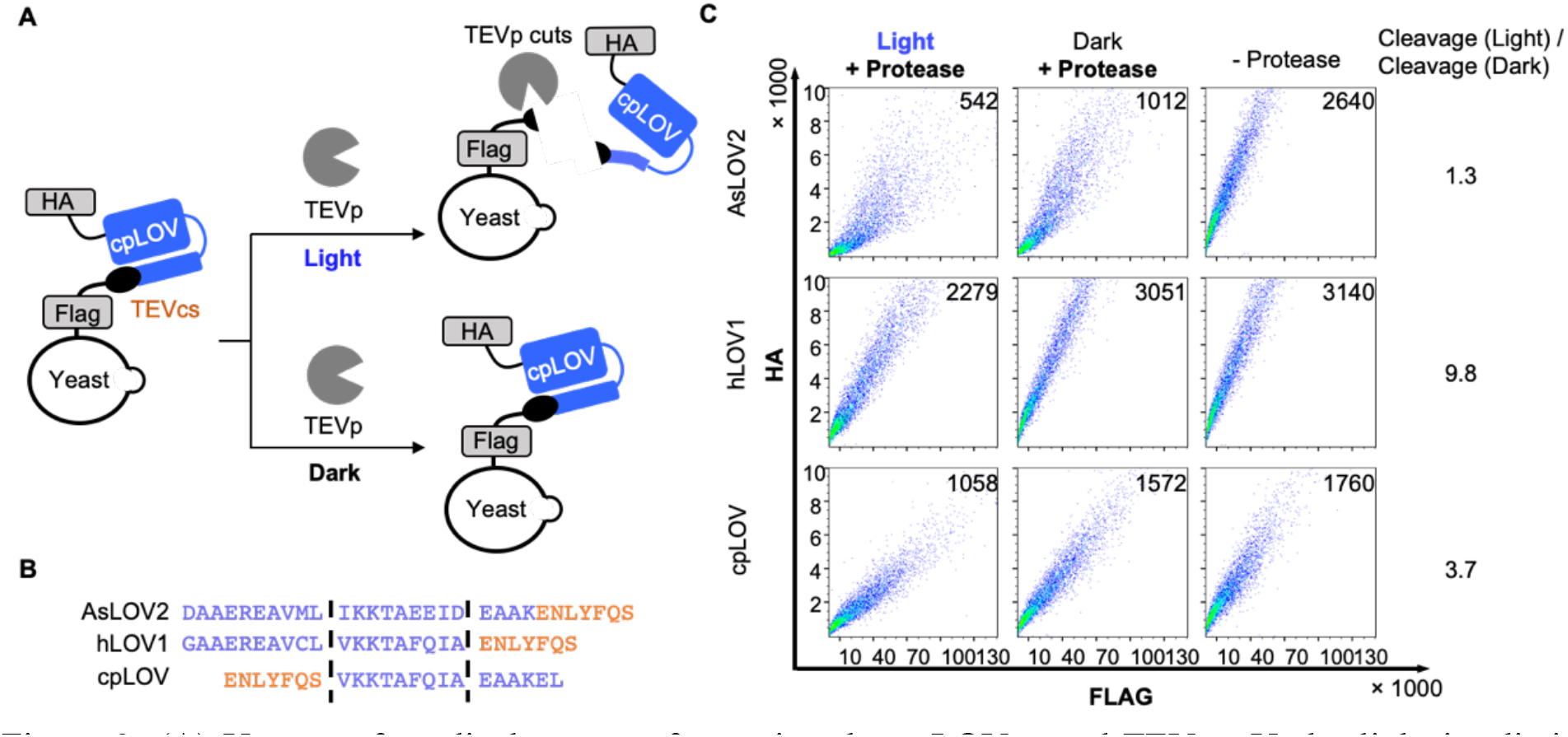
(A) Yeast surface display assay for testing the cpLOV caged-TEVcs. Under light irradiation, TEVcs is uncaged and cleaved by TEVp, causing a reduction of HA signal. (B) The Jα helix sequences and truncation sites for AsLOV2, hLOV, and cpLOV. TEVcs sequence, ENLYFQS. (C) Flow cytometry analysis of different LOV domain-caged TEVcs. The value within the plot indicates the median HA signal. Cleavage (light) / Cleavage (dark) is defined as the ratio of the difference between the median HA signal of the Light + Protease and – Protease to that of the Dark + protease and – Protease.

**Figure 4:**
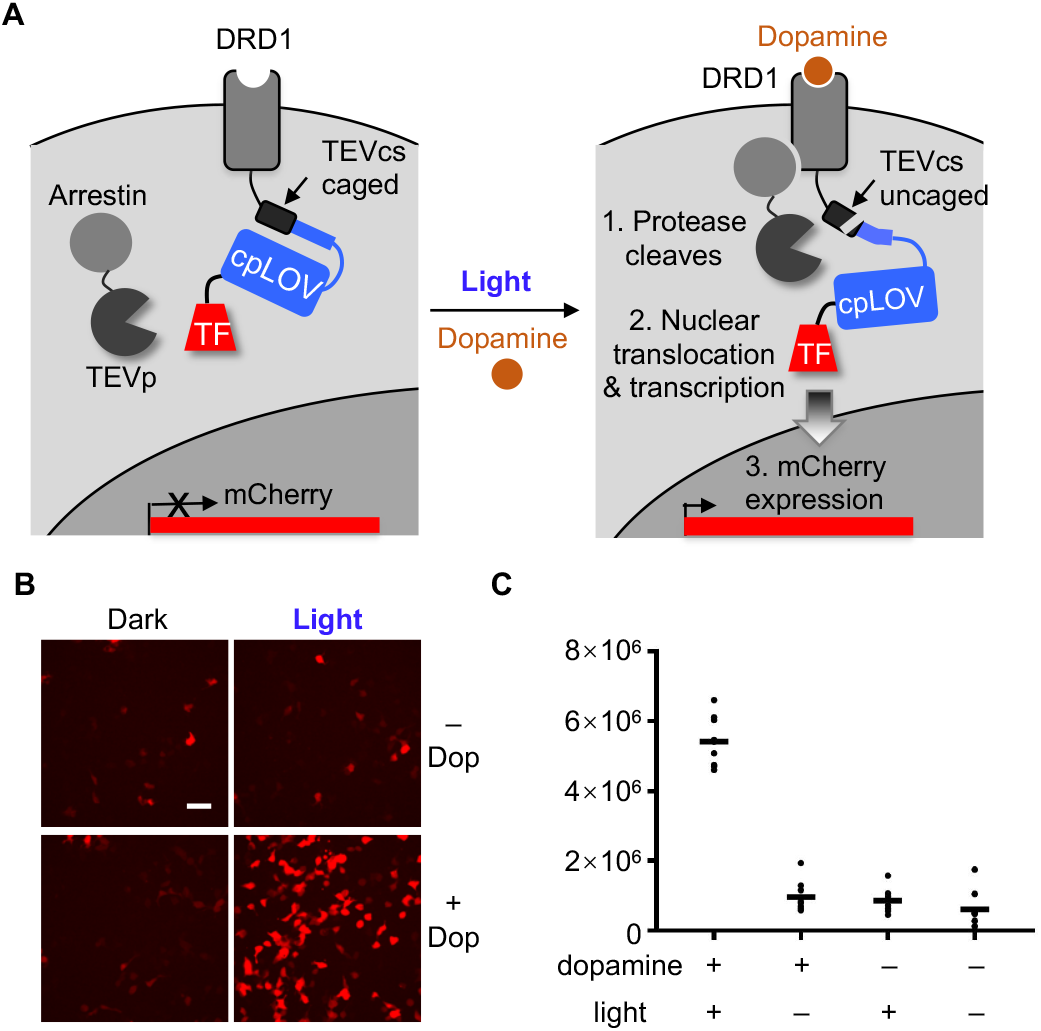
(A) Scheme of transcriptional assay in HEK293T cells. Under dark and no dopamine condition, the TEVcs is caged and not cleaved. Hence, no mCherry reporter gene is expressed. Under light and dopamine stimulation, the TEVcs is uncaged and cleaved by the protease brought into its proximity via the DRD1-arrestion interaction. The transcription factor is released from the membrane to initiate reporter gene expression. (B) Confocal fluorescence images of the transcriptional assay in (A). The cells were stimulated with light and/or 100 μM dopamine for 10 minutes. Scale bar, 50 μm. (C) Quantification of the mCherry reporter gene expression in (B).

We next investigated the effects of dual caging in improving the peptide caging in the dark. One of the long-standing challenges associated with using light-sensing proteins is the insufficient caging and the consequent high background activity in the dark state^3^. Although extensive engineering efforts have improved the dark-state caging of AsLOV2 via rational mutagenesis^6^ and directed evolution^15,30^, a general approach to improve the peptide caging efficiency in the dark will be highly advantageous. Tighter caging in the dark is especially important for experiments that are highly sensitive or require expression of the protein for an extended period of time. We performed a proof-of-concept study of dual caging of the TEVcs peptide with hLOV1 and cpLOV. We used the highly sensitive transcriptional assay similar to Fig. 4 except that the TEVcs is caged by both hLOV1 and cpLOV (Fig. 5A). During a 72-hour lentiviral transduction period, dual caging reduced the background by over 90% and effectively shifted the dynamic range to the lower end (Fig. 5B and 5C).

**Figure 5:**
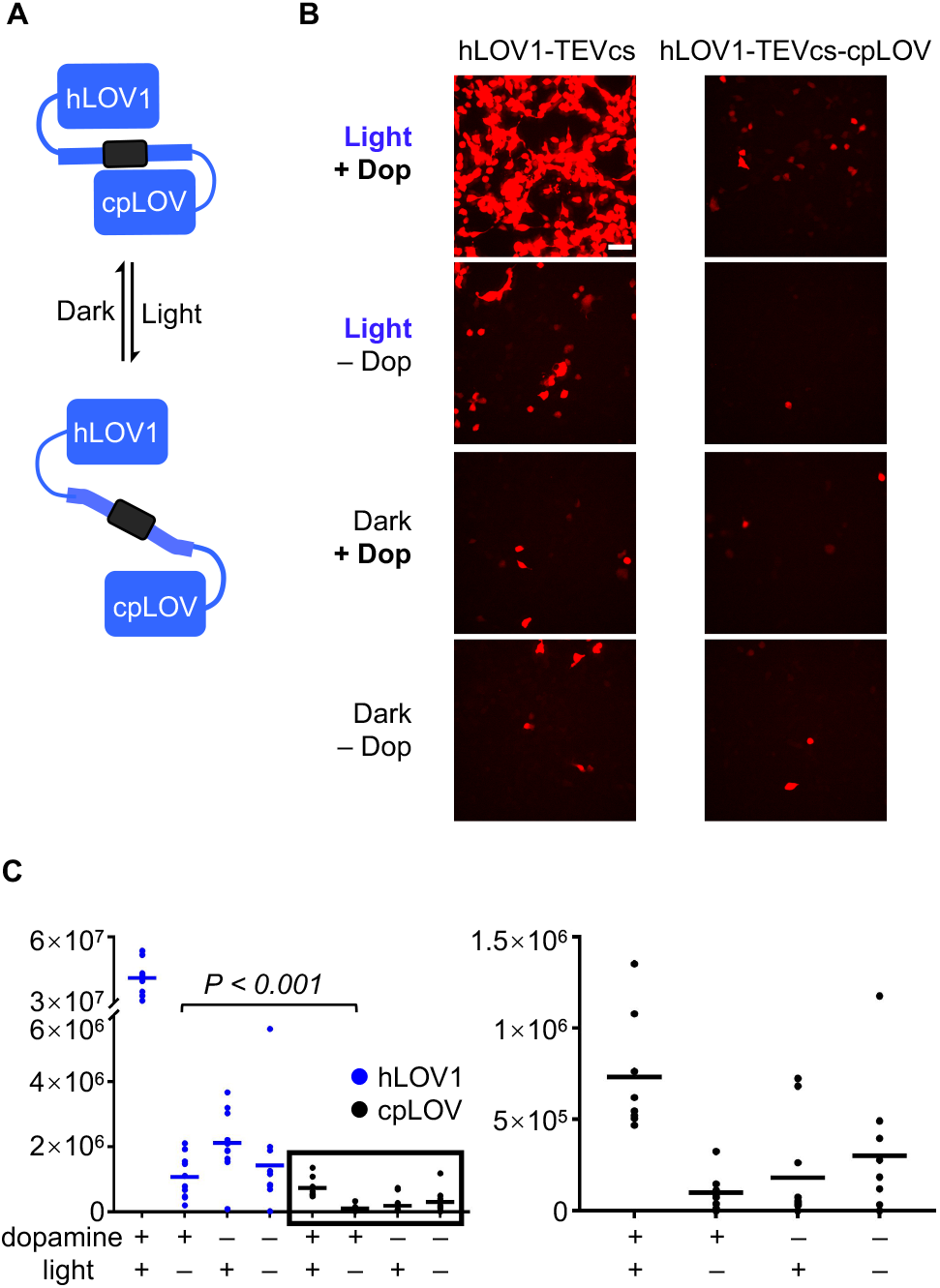
(A) Dual-caging of TEVcs. (B) Confocal fluorescence images of the dual-caged-TEVcs transcriptional assay in HEK293T cells. The cells were stimulated with light and/or 100 μM dopamine 30 minutes. Scale bar, 50 μm. C) Quantification of the mCherry reporter gene expression in (B). P value is determined by unpaired two-tailed t-test.

In conclusion, we re-engineered a light-sensing LOV domain to control peptides’ activities via fusion to their C-terminus. This new photoswitch, termed cpLOV, complemented the original AsLOV2 domain by enabling the caging of the C-terminus of a peptide, which is important for modulating the activity of peptides that require a free N-terminus. We also demonstrated for the first time that the dual caging strategy is effective in enhancing the dark state caging and could drastically shift the dynamic range of the optogenetic tool. Previous studies showed that the hinge region connecting the Jα-helix and the LOV core domain is critical for engineering a high light-dependence^15^. This is because the residues at the hinge region could introduce new interactions with the protein core to stabilize the dark state Jα-helix packing against the core. While the current version of cpLOV shows lower light-dependence than the hLOV1 domain, further engineering of the four-amino-acid glycine- and serine-linker in cpLOV can potentially improve the cpLOV’s light-dependence. The cpLOV’s light dependence can also be further improved via yeast surface display platform-based directed evolution methods in the future. Overall, cpLOV will significantly increase the application scope of light sensing proteins.

This work was supported by the University of Michigan.

