## supplementary information for "Circularly permuted LOV domain as an engineering module for optogenetic tools"

### Figures

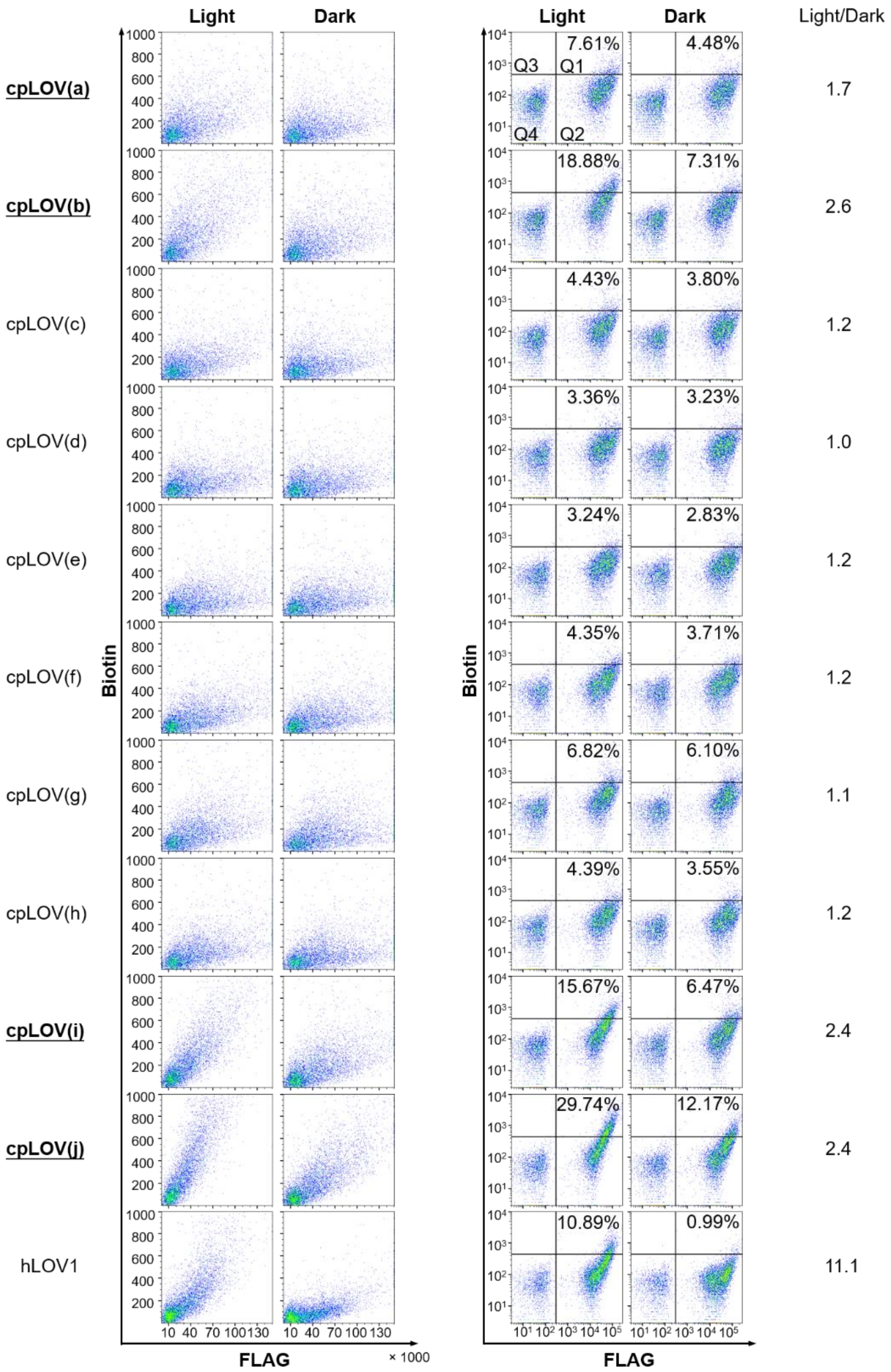


**Supplementary Fig. 1.** Related to Fig. 2E. Full sets of linear and log scale plots of the flow cytometry analysis of different SsrA-cpLOV fusion constructs. The quadrant was drawn based on the negative population in the bottom left corner in Q4. The percentage in Q1 indicates the ratio of the cell count in Q1 to that in (Q1 + Q2). The Light/Dark value is the ratio of the percentage of the light to that of the dark state. The cpLOV fusion constructs with the highest light dependence were bolded and underlined.

### Methods

**Cloning.** Yeast surface display constructs were cloned into the pCTCON2 vector. Constructs for HEK293T cell experiments were cloned into the pLX208 lentiviral vector for lentiviruses production. Constructs for protein expression and purification in *E. coli* were cloned into the pYFJ16 vector.

For cloning, PCR fragments were amplified using Q5 DNA polymerase (New England Biolabs (NEB)). The vectors were double-digested with restriction enzymes (NEB), gel purified, and ligated with gel-purified PCR fragments using T4 ligase (NEB), and Gibson assembly. Ligated DNA were heat-transformed into competent XL1-Blue *E. coli* cells.

**Expression and purification of TEV protease.** Full-length TEV protease (TEVp, S219V) was expressed as a fusion to maltose binding protein (MBP) with a polyhistidine-tag. His-tag-MBP-TEVp(S219V) transformed cells were cultured in 5 mL Miller's lysogeny broth (LB) medium (Bio Basic) supplemented with 100 mg/L ampicillin at 37 °C with shaking at 220 r.p.m. overnight for 12 h. This saturated culture was transferred to 500 mL LB with 100 mg/L ampicillin and grown at 37 °C with shaking at 220 r.p.m. for 2-3 h until $OD_{600}$ reaches 0.4-0.8. Isopropyl β-D-1-thiogalactopyranoside (IPTG, MilliporeSigma) was added to the culture to a final concentration of 1 mM for protein expression induction. The culture was grown at 16 °C with shaking at 220 r.p.m. for additional 16-24 h. Cells were then harvested by centrifugation at 4,000 × *g* for 5 min. The cell pellet was resuspended with 15 mL ice-cold B-PER (Thermo Fisher Scientific) supplemented with 1 mM Dithiothreitol (DTT, Thermo Fisher Scientific) and 100 units/mL benzonase nuclease (MilliporeSigma). The mixture was incubated on ice for 5-10 min, and centrifuged at 17,000 × *g* for 15 min to clarify the cell lysate. The clarified cell lysate was incubated with 3 mL Ni-NTA resin slurry (Thermo Fisher Scientific) for 10 min with rotation and then transferred to a gravity column. The resin was washed with 5 mL washing buffer (30 mM imidazole, 50 mM Tris, 300 mM sodium chloride, 1 mM DTT, pH = 7.8). The protein was eluted with 3 mL elution buffer (200 mM imidazole, 50 mM Tris, 300 mM sodium chloride, 1 mM DTT, pH = 7.8). The eluent was concentrated to about 1/10 of its original volume with a 15 mL 10,000 Da cutoff centrifugal unit (MilliporeSigma). The concentrated proteins were mixed with 80% (v/v) glycerol to a final concentration of 30% and then flash frozen in liquid nitrogen and stored at -80 °C.

**Expression and purification of SspB-APEX2.** SspB-APEX2 was expressed with polyhistidine-tag in BL21 *E. coli* cells following the same procedures as "Expression and purification of TEV protease" except that DTT was not added into the cell lysate, washing buffer, or elution buffer.

**Yeast culture.** Yeast display plasmids in pCTCON2 were transformed into *Saccharomyces cerevisiae* strain EBY100 competent cells following the kit protocol by Zymo Research. Briefly, 1 µg of the DNA was mixed with 5 µL yeast competent cells, and 200 µL of Frozen-EZ Yeast Solution 3 (Zymo Research). The cells were incubated at 30 °C for 30 min to 2 h, and then transferred to 5 mL synthetic dextrose plus casein amino acid media (SDCAA, 20 g/L dextrose, 6.7 g/L yeast nitrogen base without amino acids (BD Difco), 5 g/L casamino acids (BD Difco), 5.4 g/L disodium phosphate, 8.56 g/L monosodium phosphate in deionized water). The cells were grown at 30 °C with shaking at 220 r.p.m. After the initial saturation ($OD_{600}$ > 10) in 2-3 days, the yeast culture was passaged at least once in SDCAA prior to protein expression induction. To induce expression of the pCTCON2 plasmid, 500 µL of the overnight yeast in SDCAA media was added to 5 mL of SGCAA (synthetic galactose plus casein amino acid media, 20 g/L galactose, 6.7 g/L yeast nitrogen base without amino acids (BD Difco), 5 g/L Casamino acids (BD Difco), 5.4 g/L Disodium phosphate, 8.56 g/L Monosodium phosphate)) media and let grow at 30 °C with shaking at 220 r.p.m. overnight.

**Yeast labeling.** 250 µL of yeast cells induced in SGCAA overnight were mixed with 1 mL PBSB (sterile phosphate-buffered saline supplemented with 1 g/L bovine serum albumin), and then centrifuged at 6,000 × *g* for 30 s. The cell pellet was resuspended and washed once with 1 mL PBSB.

Yeast expressing SsrA was subject to APEX2 labeling[1] prior to antibody labeling. For the light condition, the yeast samples were irradiated with blue LED light all the time except during centrifugation before antibody labeling. For the dark condition, experiments are performed in a dark room with a red light. Yeast samples were incubated with 100 µL of SspB-APEX2 solution (expressed and purified as described above, under "Expression and purification of SspB-APEX2") in the light or in the dark at room temperature for 10 min with rotation. After incubation, samples were washed twice with 1 mL PBSB, and further resuspended in 950 µL PBSB with 1% bovine serum albumin (BSA). 1 µL biotin-phenol (1 mM in dimethyl sulfoxide) was added and thoroughly mixed with the sample by vortexing. Then, 1 µL of hydrogen peroxide (0.5 mM in water, freshly prepared, MilliporeSigma) was added and thoroughly mixed by vortexing. After exactly 2 min, 200 µL of quenching solution 1 (30 mM Trolox (Thermo Fisher Scientific), 60 mM sodium ascorbate (MilliporeSigma), freshly prepared) was added to quench the reaction. The samples were centrifuged at 6,000 × *g* for 30 s, and the supernatant was discarded. 400 µL of quenching solution 2 (5 mM Trolox, 10 mM sodium ascorbate, freshly prepared) was then added. After another centrifugation at 6,000 × *g* for 30 s, the supernatant was discarded, and the sample was washed twice with 1 mL PBSB. Samples were then incubated with 100uL PBSB solution with mouse anti-FLAG antibody (2.5 µM, Sigma) and streptavidin-phycoerythrin (PE) (200-fold dilution, Jackson ImmunoResearch) at room temperature for 15 min with rotation, washed twice with 1 mL PBSB, and incubated with Alexa Fluor 647 anti-mouse antibody at 2.5 µM in 100 µL PBSB) at room temperature for 15 min with rotation. Samples were washed twice and resuspended in 1 mL PBSB for FACS within 24 h as described under "FACS analysis and library selection".

Yeast cells that expressed TEV protease cleavage site (TEVcs) was first treated with TEVp, then labeled with antibody-fluorophore conjugate. Samples were incubated with 200 µL of PBSB containing TEVp (expressed and purified as described above, under "Expression and purification of TEV protease") in the light or in the dark for 3 h with rotation. For negative control, TEVp were omitted. 30 mM reduced and 3 mM oxidized glutathione (MilliporeSigma) were added to all samples to keep TEVp under reducing conditions. After incubation with TEVp, samples were washed twice with PBSB and subsequently incubated with primary antibodies (mouse anti-FLAG and rabbit anti-HA antibodies at 2.5 µM each in 100 µL PBSB) and secondary antibodies (Alexa Fluor 647 anti-mouse and Alexa Fluor 568 anti-rabbit antibodies at 2.5 µM each in 100 µL PBSB) at room temperature for 15 min with rotation. Two washes with PBSB were performed after each step of labeling. Samples were resuspended in PBSB for FACS within 24 h as described under "FACS analysis and library selection".

**FACS analysis.** Labeled yeast cells were analyzed with an LSRFortessa cell analyzer flow cytometer (BD Biosciences) equipped with 640 nm laser and 670/14 nm emission filter (for Alexa Fluor 647) as well as 561 nm laser and 586/15 nm emission filter (for Alexa Fluor 568 and PE). Library samples were sorted with a FACSAria III cell sorter flow cytometer (BD Biosciences) equipped with 633 nm laser and 660/20 nm emission filter (for Alexa Fluor 647) as well as 561 nm laser and 582/15 nm emission filter (for Alexa Fluor 568 and PE).

FACS data are analyzed by FlowJo. For cpLOV caging SsrA experiment, the quad was gated according to the Flag signal negative cell population on the log scale plot. The percentage is calculated by Q1/(Q1 + Q2), where Q1 and Q2 stands for the cell number in the quad respectively. For cpLOV caging TEVcs experiment, a quad was gated according to the Flag signal negative cell population on the log scale plot. And then the median for Flag signal positive cells were taken. Cleavage (Light) = median(-Protease) – median(Light, +Protease). Cleavage (Dark) = median(-Protease) – median(Dark, +Protease).

**HEK 293T cell culture.** HEK293T cells less than 20 passages were cultured at 37 °C under 5% $CO_2$ in T25 or T75 flasks in complete growth media (1:1 Dulbecco's Modified Eagle Medium (DMEM, Gibco): Minimum Essential Media (MEM, Gibco) supplemented with 10% fetal bovine serum (FBS, by volume, MilliporeSigma), 50 mM HEPES (Gibco), 50 units/mL penicillin, and 50 µg/mL streptomycin).

**Production of lentivirus supernatant for HEK 293T cell transduction.** New cell culture flasks were incubated with 20 µg/mL human fibronectin (HFN, MilliporeSigma) at 37 °C for at least 10 min. After incubation, HFN was aspirated, and HEK293T cells (less than 20 passages) were plated at 70-90% confluence. For a T25 flask, cells were incubated at 37 °C for 1-3 h. 2.5 µg viral DNA, 0.25 µg pVSVG, and 2.25 µg delta8.9 lentiviral helper plasmid were mixed and diluted with 250 µL of DMEM. Then, 25 µL PEI MAX solution was added to the DNA mixture. The mixture was incubated at room temperature for at least 10 min, mixed with 1 mL complete growth media, and transferred to the T25 flask. Cells were incubated at 37 °C for 48 h, and the supernatant solution was collected, flash frozen in liquid nitrogen, and stored at -80 °C before use.

**HEK 293T cell lentiviral transduction for transcriptional assay.** HEK 293T cells less than 20 passages were cultured at 37 °C under 5% $CO_2$ in T25 or T75 flasks in complete growth media. 48-well plates were pretreated with 200 µL 20 µg/mL HFN for 10 min at 37 °C. HEK 293T cells were then plated at 40%-60% confluence. For transduction of a single well in a 24-well plate, 100 µL of each supernatant virus (UAS-mCherry reporter gene, DRD1-TEVcs-cpLOV-TF, Arrestin-TEVp) was added gently to the top of the media. The plates were wrapped with aluminum foil and incubated at 37 °C under 5% $CO_2$ for 48 h before stimulation. The culture media was aspirated and the cells were stimulated with 200 µL 100 µM dopamine hydrochloride (Alfa Aesar) in complete growth media or white light for 10 or 30 min. The dopamine solution was then aspirated and the cells were washed with 200 µL complete growth media for three times. For the cells that were not treated with dopamine, complete growth media was used in stimulation step. The plates were wrapped with aluminum foil and incubated at 37 °C under 5% $CO_2$ for 24 h (cpLOV single caged TEVcs in Fig. 4) or 48 h (dual caged TEVcs in Fig. 5) before fluorescence microscope imaging.

**Fluorescence microscopy of cultured cells.** Confocal imaging was performed on a Nikon inverted confocal microscope with 10× air, 20× air, and 60× oil-immersion objectives, outfitted with a Yokogawa CSU-X1 5000RPM spinning disk confocal head, and Ti2-ND-P perfect focus system 4, a compact 4-line laser source: 405 nm (100 mW), 488 nm (100 mW), 561 nm (100 mW) and 640 nm (75 mW) lasers. The following combinations of laser excitation and emission filters were used for various fluorophores: DAPI (405 nm excitation; 455/50 nm emission), EGFP/Alexa Fluor 488 (488 nm excitation; 525/36 nm emission), mCherry/Alexa Fluor 568 (568 nm excitation; 605/52 nm emission), Alexa Fluor 647 (647 nm excitation; 705/72 nm emission). ORCA-Flash 4.0 LT+sCMOS camera. Acquisition times ranged from 100 to 1000 msec. All images were collected and processed using Nikon NIS-Elements hardware control and analysis module.

**Fluorescence microscopy image analysis** For confocal images, 10-12 fields of view per well were taken. Confocal fluorescence microscopy images were analyzed using the General Analysis 3 module on the Nikon NIS-Elements AR Analysis software. For the transcriptional assay reporter gene channel, a threshold was set to be just above the background fluorescence. Signal above the threshold from each field of view was integrated and plotted as dot plots using GraphPad Prism 8. Mean intensity was calculated for each condition.
